# Attenuation hotspots in neurotropic human astroviruses

**DOI:** 10.1101/2022.09.06.506719

**Authors:** Hashim Ali, Aleksei Lulla, Alex S. Nicholson, Elizabeth B. Wignall-Fleming, Rhian L. O’Connor, Diem-Lan Vu, Stephen C. Graham, Janet E. Deane, Susana Guix, Valeria Lulla

## Abstract

During the last decade, the detection of neurotropic astroviruses has increased dramatically. The MLB genogroup of astroviruses represents a genetically distinct group of zoonotic astroviruses associated with gastroenteritis and severe neurological complications in young children, the immunocompromised and the elderly. Using different virus evolution approaches, we identified dispensable regions in the 3′ end of the capsid-coding region responsible for attenuation of MLB astroviruses in susceptible cell lines. To create recombinant viruses with identified deletions, MLB reverse genetics and replicon systems were developed. Recombinant truncated MLB viruses had wild type-like or enhanced growth and replication properties in permissive cells but were strongly attenuated in iPSC-derived neuronal cultures confirming the location of neurotropism determinants. This approach can be used for the development of vaccine candidates using attenuated astroviruses that infect humans, livestock animals and poultry.

## Introduction

Human astroviruses (HAstVs) belong to the genus *Mamastrovirus*, family *Astroviridae* and are a common cause of gastroenteritis in children, the elderly and immunocompromised adults ^1^. Lately, the HAstV group of the *Astroviridae* family has expanded to include new groups of viruses unrelated to the eight previously described classic HAstV serotypes (Fig. 1A). These new human astrovirus groups are more closely related to certain animal astroviruses than to the classical HAstVs, suggesting zoonotic transmission ^1^. One of these groups is named MLB, after the first novel human astrovirus described in 2008 in Melbourne (Australia) identified in feces of pediatric patients with gastroenteritis. Later, the MLB group of HAstVs was assigned to a neurovirulent group of astroviruses due to the association with severe cases of meningitis/encephalitis, febrile illness, and respiratory syndromes ^2^. Interestingly, it was recently shown that astroviruses found in the fecal samples of macaque monkeys were genetically similar to human astrovirus MLB and caused chronic diarrhea ^3^.

**Figure 1.**
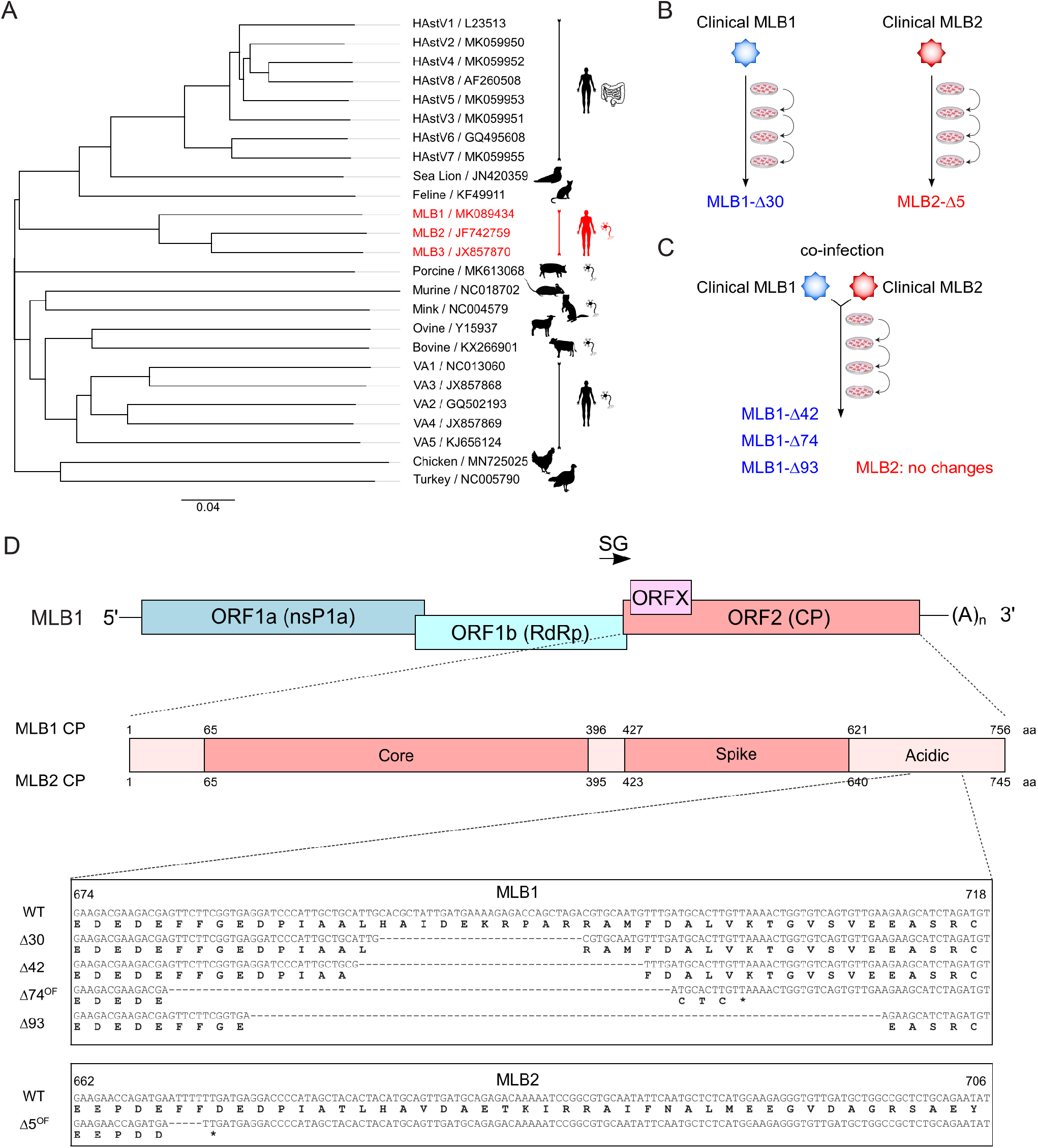
Classification and attenuation of MLB astroviruses. **(A)** Simplified phylogenetic tree for the *Astrovirus* genus. The tree is based on full nucleotide sequences available for indicated species. The pictogram of intestine (HAstV1-8) or neuron (several species) indicates the tropism associated with astrovirus strains. Neurotropic MLB strains are shown in red. **(B)** The evolution experiment was performed for MLB1 and MLB2 astroviruses. **(C)** The co-evolution experiment was performed for MLB1 and MLB2 astroviruses. **(D)** Nucleotide and amino acid sequences of MLB1 and MLB2 viruses containing deletions identified in evolved MLB virus stocks.

HAstVs are small, non-enveloped, icosahedral viruses with positive-sense single-stranded RNA genome containing 5′ untranslated region (UTR), four open reading frames (ORF1a, ORF1b, ORFX and ORF2) and a 3′ UTR with poly A tail ^4,5^. ORF1a encodes non-structural polyprotein nsP1a, ORF1b is expressed via ribosomal frameshifting mechanism and encodes RNA-dependent RNA polymerase (RdRp). The subgenomic (SG) RNA encodes two ORFs – ORF2 and ORFX, the latter encoding a viroporin ^5^. The product of the ORF2 coding sequence is translated into the structural capsid polyprotein (CP) of about 72-90 kDa, depending on the virus strain ^6^, which then undergoes C-terminal cleavage by cellular caspases ^7^. Despite the required function of caspases, the astrovirus release is described as an unclassified nonlytic process ^8^. In some astroviruses, the structural polyprotein is cleaved by trypsin resulting in the formation of 3′ truncated (25-34 kDa) proteins. Maturation of the astrovirus capsid protein is a very dynamic process, transforming the virus from a non-infectious intracellular form (VP90) to a primed extracellular form (VP70), and finally generating an icosahedral infectious mature virion (VP34/27/25). However, some concerns remain to be addressed to fully understand the astroviruses capsid assembly and maturation. In particular, what differences in the capsid protein of the MLB genotypes make them different from the classical astroviruses and what determines their infectivity? It has been shown that trypsin treatment of classical astroviruses increases their infectivity ^9,10^, whilst not affecting the MLB genotypes ^2^. The mechanism of capsid polyprotein cleavage and the functional role of cleaved CP in the MLB group of astroviruses is not yet understood. It is also unclear if MLB astroviruses exploit cellular proteases other than trypsin to process the capsid protein and how this impacts the infectivity of virus particles.

Growing evidence suggests that astroviruses are found globally, infecting a wide range of species, and have the potential for recombination, rapid evolution, and can adapt to different hosts ^3,11–16^. Unfortunately, many astrovirus groups have remained overlooked for decades because of the absence of molecular tools, such as infectious clones and replicons. Therefore, developing a robust reverse genetics (RG) system for the non-classical human astroviruses is essential to understand the basic biology, evolution and host-virus interplay.

In 1997 Matsui’s group established the first RG system for the human astrovirus serotype 1 to rescue infectious viral particles ^17^. This system has been successfully used and has shed light on multiple aspects of astrovirus replication and pathogenesis. HAstV1 RG system requires two cell lines to recover infectious particles: the transfection of BHK-21 cells with *in vitro* transcribed viral RNA and then propagation of the obtained supernatant in the permissive Caco-2 cells in the presence of trypsin. Several DNA-based RG systems were developed, including efficient chimeric HAstV1/8 RG system ^18,19^, however, all of them relied on two cell lines and were limited to classical human astroviruses. So far, RG systems for two non-human astroviruses were developed: first for the avian astrovirus by using duck astrovirus (DAstV) genome of D51 strain ^20^, and second for porcine astrovirus (PAstV1-GX1) ^21^. Although both non-human RG systems allow the recovery of infectious viral particles, these systems also rely on the two cell lines.

It is therefore essential to develop the RG system for neurotropic astroviruses to understand the molecular determinants for neurotropism and neurovirulence. Here we report the RG system for two non-classical human neurotropic astroviruses that relies on a single cell line and can be used to rescue and propagate MLB1 and MLB2 human astroviruses. We also developed a set of detection tools as well as replicon systems for both MLB astroviruses. Using this system we identified and characterized attenuation hotspots located at the 3′ end of the MLB genomes. In the future, this RG system will deepen the understanding of molecular virology of MLB-group astroviruses and allow the design of tools to address open questions on viral evolution, replication, packaging and pathogenesis.

## Results

### Evolution and cell culture adaptation of neurotropic MLB astroviruses

The selection for attenuated viruses through serial passaging in highly susceptible cells is a well-known approach for directed evolution ^22^. We, therefore, hypothesized that this strategy could be applied to attenuate MLB astroviruses. Clinical MLB isolates were passaged in susceptible cell lines as previously described ^2^ (Fig. 1B). Sequencing of a passaged clinical MLB1 isolate revealed a deletion of 30 nucleotides in the 3′ end of the genome spanning into the coding sequence of CP. A similar region was affected in the passaged MLB2 clinical isolate – a single out-of-frame deletion of 5 nucleotides in the 3′ part of the genome (Fig. 1D).

Another strategy for directed virus evolution is based on co-infection of closely related virus species. The better replicating “partner” can either out-compete or complement the replication of another virus. To test this hypothesis, we co-infected Huh7.5.1 cells at a multiplicity of infection (MOI) 0.1 with MLB1 and MLB2 viruses (Fig. 1C). This resulted in simultaneous replication and propagation of both strains on passaging without detected out-competition or recombination for 10 consecutive passages. Interestingly, no changes were observed in MLB2 genomes; however, several in-frame and out-of-frame deletions were detected in the 3′ part of the MLB1 genome further confirming the instability of the 3′ region in this virus (Fig. 1D).

Elucidating the functional significance of the identified deletions requires MLB astrovirus detection tools and would be dramatically accelerated by the establishment of a robust RG system. We, therefore, aimed to create these essential tools.

### Cell culture models and detection tools for neurotropic MLB1 and MLB2 astroviruses

First, we developed a set of essential tools for specific immune detection of virus infection. The folded region of MLB1 capsid protein corresponding to amino acids 61-396 of ORF2-encoded polyprotein possessing a C-terminal 8×His-tag (Fig. 2A) was used for bacterial expression and affinity purification, resulting in homogeneous CP_NTD_ protein (Fig. 2B). The purified recombinant protein was used for the production of highly sensitive antibodies allowing the detection of ≤ 1 ng of the purified CP_NTD_ of MLB1 (Fig. 2C). Due to 95% identity between corresponding domains of MLB1 and MLB2 CPs, polyclonal antibodies were expected to cross-detect capsid proteins derived from both strains. Indeed, it specifically recognized capsid proteins from MLB1- and MLB2-infected cells (Fig. 2C).

**Figure 2.**
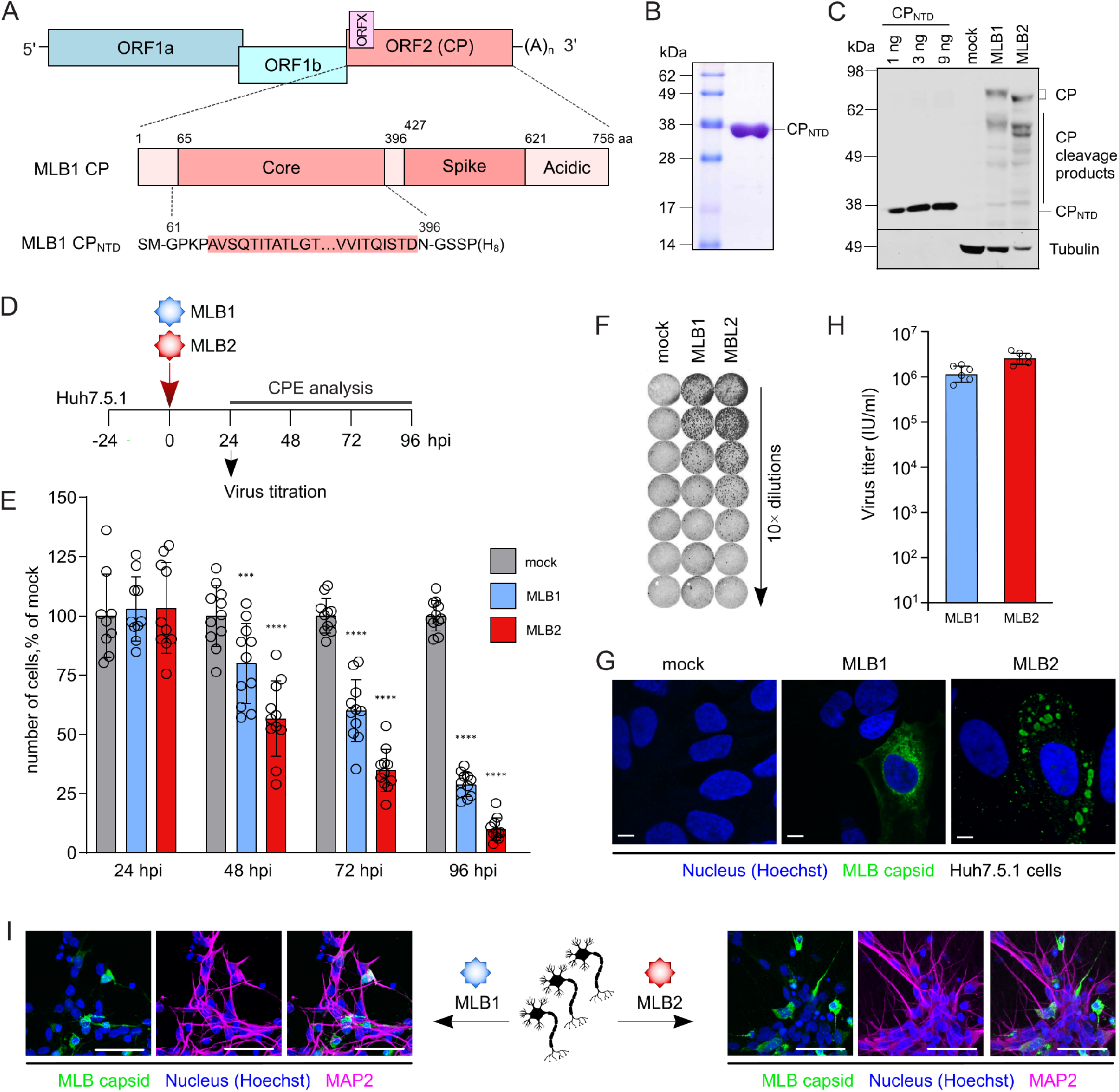
Purification of CPNTD and immunodetection of MLB1- and MLB2-infected Huh7.5.1 cells and iPSC-derived neurons. **(A)** Schematic representation of MLB1 genome and location of CP_NTD_. Lower panel represents the sequence of recombinant CP_NTD_. ORF, open reading frame; CP, capsid protein; NTD, N-terminal domain. **(B)** Coomassie-stained SDS-PAGE profile of the purified CP_NTD_ from *E. Coli*. **(C)** Huh7.5.1 cells were infected with MLB1 and MLB2 viruses at MOI 0.1 and incubated for 48 hours. CP was detected using the antibody generated against CP_NTD_. Purified CP_NTD_ was used for detection limit assessment (1-9 ng). **(D)** Experimental setup to determine the cytopathic effect (CPE) and titration for MLB1 and MLB2 infection. **(E)** Huh7.5.1 cells were infected at an MOI 1 and incubated for indicated periods, then washed with media, stained and imaged. Hoechst-stained nuclei were counted from 12 images (∼200 cells per image) and normalized to mock-infected samples. Data are mean ± SEM. ***p < 0.001, ****p < 0.0001 using two-way ANOVA test against mock. **(F)** Huh7.5.1 cells were seeded on 96-well plate and infected with 10-fold serial dilutions of MLB1 and MLB2 astroviruses, fixed at 20-24 hpi, permeabilized, stained with anti-CP antibody, and imaged by LI-COR. **(G)** Huh7.5.1 cells were infected with MLB1 and MLB2 viruses and incubated for 24 h. Representative confocal images of fixed and permeabilized cells visualized for CP (green) and stained for nuclei (Hoechst, blue) are shown. Scale bars are 10 µm. **(H)** Huh7.5.1 cells were infected with MLB1 and MLB2 virus stocks at MOI 0.1 and incubated for 72-120 hours. Virus titers were determined from 6 independent experiments. Data are mean ± SEM. **(I)** i^3^Neurons were seeded on IBIDI plates, differentiated into mature glutamatergic neurons, infected with MLB1 and MLB2 viruses and incubated for 48 (MLB2) or 96 (MLB1) hours. Representative confocal images of fixed and permeabilized cells visualized for MLB CP (green), neuronal marker MAP2 (magenta) and stained for nuclei (Hoechst, blue) are shown. Scale bars are 50 µm.

When passaging clinical isolates, we noticed that the Huh7.5.1 cell line supports MLB replication and results in a moderate cytopathic effect (CPE). MLB clade viruses were previously reported to replicate in Huh7 and Huh7.5 cell lines with the ability to establish a persistent infection on passaging ^2^. Presumably, in contrast to the immune-competent Huh7 cells, the more susceptible Huh7.5.1 cell line ^5^ can allow for enhanced replication and development of CPE. This cell line was also reported to support active replication of the classical human astrovirus 1 (HAstV1) ^23^. The CPE was apparent for MLB2 at 24-48 hours post infection, whereas slower replicating MLB1 showed CPE at 48-72 hours post infection, reaching >60% cell death at 3 days post infection for MLB2 and at 4 days post infection for MLB1 (Fig. 2D-E).

The permissiveness of the Huh7.5.1 cell line allowed the development of a virus titration system. Cells cultured on 96-well plates were infected with 10-fold dilutions of MLB1 and MLB2 stocks, fixed and stained with CP_NTD_-antibody using in-cell near-infrared fluorescence-based detection (Fig. 2D, F). The cytoplasmic distribution of CP was confirmed by confocal microscopy further demonstrating that both MLB1 and MLB2 can be detected 24 hours post infection (Fig. 2G). Since no released virus could be detected after 20-24 hpi, this timepoint was utilized for single-round infection experiments like titration of the virus stocks. Efficient virus release was detected at 48-72 hpi for MLB2 and at 96-120 hpi for MLB1, reaching 1-3×10^6^ infectious units (IU) per ml (Fig. 2H).

Finally, to confirm the neurotropic properties of MLB astroviruses, we developed a physiologically relevant system to infect and monitor MLB infection in neurons. The neurotropism of MLB astroviruses was previously described and indicates their ability to infect cells of neuronal origin ^2,24^. The recently developed methodology to efficiently differentiate human induced pluripotent stem cells (iPSCs) into isogenic cortical glutamatergic neurons (i^3^Neurons) ^25^ provided a suitable platform to experimentally assess neurotropic properties of MLB1 and MLB2 viruses. Both viruses resulted in efficient infection at 48 hpi (MLB2) and 96 hpi (MLB1) further confirming the ability of these viruses to infect post-mitotic neuronal cells (Fig. 2I).

### Development, annotation and assessment of infectious clones for MLB1 and MLB2 viruses

The 5′ and 3′ terminal consensus sequences were used to design specific primers to amplify MLB1 (MK089434) and MLB2 (JF742759) full-length genomes. The entire genomes of MLB1 (Fig. 3A) and MLB2 (Fig. 3B) were cloned into the T7 promoter-containing plasmid using a single-step ligation independent cloning. The obtained plasmids were sequenced and the ORFs and functional elements were annotated based on homology with other astroviruses (GenBank accession numbers: ON398705, ON398706). The resulting differences between recombinant and clinical isolate sequences have arisen due to polymorphisms present in initial clinical isolates. To produce recombinant viruses, the infectious clones of MLB1 and MLB2 were linearized and full genomic RNAs were synthesized *in vitro* using T7 RNA polymerase. Huh7.5.1 cells were electroporated with transcribed RNAs and incubated for 48 hours (MLB2) or 72 hours (MLB1) until the appearance of CPE. The supernatants were titrated and passaged at MOI 0.1 followed by titration and sequencing of resulting viral genomes (Fig. 3C). The obtained recombinant viruses recapitulated the growth properties of the original clinically isolated MLB astroviruses ^2^, reaching final titers of 10^6^-10^7^ IU/ml. Consistent with previous findings ^2^, the increase of the virus in the extracellular fraction was higher for MLB2 than for MLB1 (Fig. 3D-E). The infection process coincided with the accumulation of capsid protein, corresponding to observed increased virus production in these samples (Fig. 3F). To get an estimation of the nature of smaller CP products, CP-specific products derived from cellular and media samples were analyzed. Interestingly, we predominantly observed cleaved CP form in the media-derived samples suggesting possible extracellular cleavage (Fig. 3G) in analogy to classical HAstV strains ^26^. Passaging of the MLB1 and MLB2 viruses resulted in similar virus titers (Fig. 3H-I) and consistent production of capsid proteins (Fig. 3J-K).

**Figure 3.**
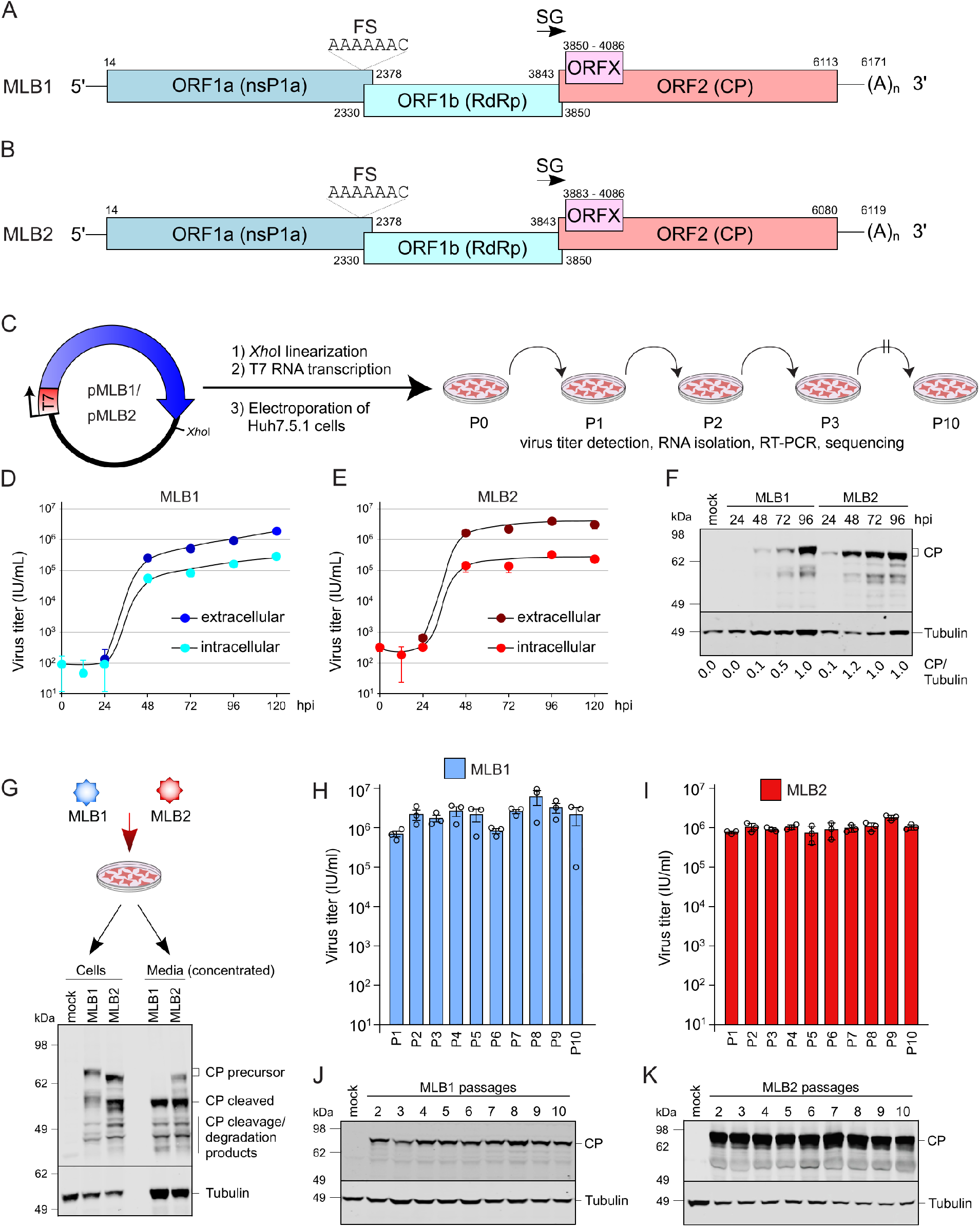
Generation and validation of reverse genetics system for MLB1 and MLB2 astroviruses. **(A-B)** Schematic representation of MLB1 (A) and MLB2 (B) genomes used to generate infectious clones. MLB genome elements: ORF, open reading frame; RdRp, RNA-dependent RNA polymerase; CP, capsid protein; FS, frameshift site; SG, subgenomic promoter. **(C)** Strategy for a plasmid-derived reverse-genetics system for MLB1 and MLB2. MLB cDNAs contain the entire genome flanked by the T7 promoter and *Xho*I linearization site. Huh7.5.1 cells were electroporated with full-genome T7 transcripts, the collected virus was used for serial passages in the same cell line. **(D-E)** Multistep growth curves of MLB1 (D) and MLB2 (E) on Huh7.5.1 cells. Cells were infected at an MOI 0.1, and virus titer was measured from the intracellular and extracellular fractions in triplicates. Data are mean ± SEM. **(F)** Cells were infected at an MOI 0.1, harvested at 48 hpi and analyzed by western blotting with anti-CP and anti-tubulin antibodies. **(G)** Huh7.5.1 cells were infected with MLB1 and MLB2 at an MOI 0.1 and incubated for 72 hours. The cell- and media-derived samples were harvested and analyzed by western blotting. **(H-I)** Huh7.5.1 cells were infected in triplicates with MLB1 (H) and MLB2 (I) at an MOI 0.1 and incubated for 72-120 hours until full CPE. Total virus titers of 10 serial passages were determined in triplicates (n=2 independent experiments). Data are mean ± SEM. **(J-K)** Analysis of CP expression in Huh7.5.1 cells infected with 2-10 passage of MLB1 (J) and MLB2 (K). Cells were infected at an MOI 0.1, harvested at 48 hpi and analyzed by western blotting.

### Assessing genome stability of MLB1 and MLB2 viruses

To evaluate the MLBs genome stabilities, we performed serial passaging by using RG-derived MLB1 and MLB2 recombinant viruses. The passaging was performed in biological duplicate, starting from *in vitro* RNA transcripts. No changes were detected in the passaged recombinant MLB2 virus, suggesting that its genome is stable in Huh7.5.1 cells. The slower replicating recombinant MLB1 was less stable and accumulated mutations in the 3′ part of the genome. An out-of-frame single-nucleotide insertion was detected at passage 3, that co-existed with wild-type (wt) MLB1 resulting in continuous co-infection for 7 consecutive passages (Table 1). A distinct cluster of mutations at the C-terminal end of CP was identified in the second experiment (Table 1) suggesting instability of RG MLB1 genomes during longer virus passaging.

**Table 1.** Genome changes in evolved recombinant MLB1 and MLB2 astroviruses.

| Virus | Mutations (position in the genome) |
| --- | --- |
| Recombinant MLB1 | Experiment 1:<br>• insertion of A (6017) at passage 3, premature termination of CP at 729 aa, co-exist with wt for the next 7 passages<br>Experiment 2:<br>• C5059T (P406L in CP) at passage 10<br>Cluster of mutations in CP: A6030G (K730E), AA6035GG (R732G), A6046G (Y735C) at passages 2-10 |
| Recombinant MLB2 | no changes detected |

To put these mutations and previously identified deletions (Fig. 1D) in the context of naturally occurring changes in MLB genomes, the analysis of all related WT MLB astrovirus sequences was performed. As expected, the 3′ region of the MLB2 genome was very conserved: few amino acid variations and no deletions were found in publicly available MLB2 genome sequences (Fig. 4A). In contrast, the 3′ region of MLB1 was more diverse, containing multiple changes throughout the analyzed region as well as one amino acid deletion upstream of the experimentally observed deletion region (Fig. 4B), consistent with the mutation- and deletion-prone nature of the 3′ region of MLB1 genome.

**Figure 4.**
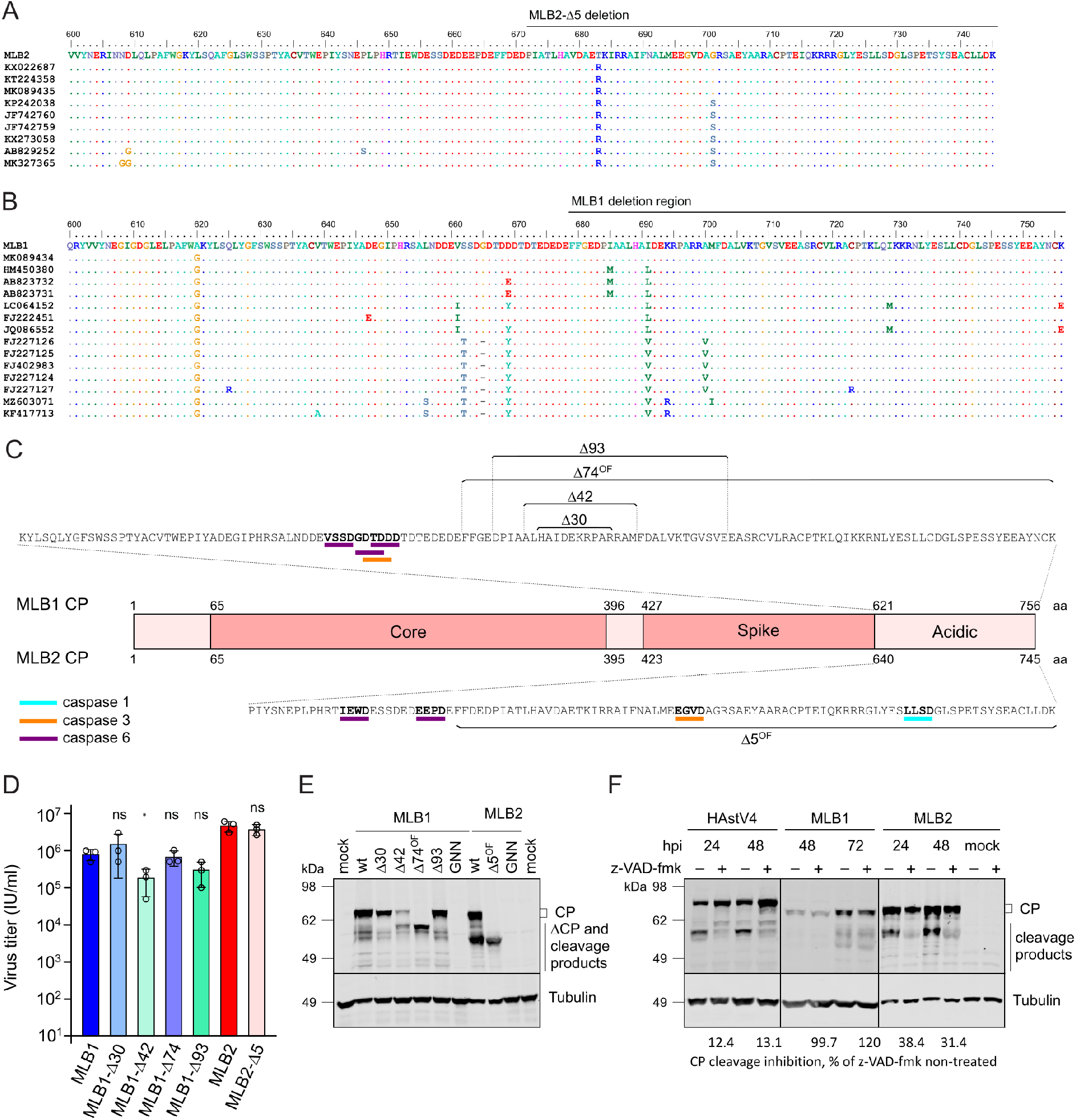
Analysis of C-terminal part of MLB1 and MLB2 astrovirus capsid polyprotein. **(A)** Analysis of publicly available sequences of C-terminal part of MLB2 CP. **(B)** Analysis of publicly available sequences of C-terminal part of MLB1 CP. The deletion region is indicated on top of MLB1 and MLB2 alignments (A-B). **(C)** Analysis of the C-terminal part of MLB1 and MLB2 CP for putative caspase cleavage sites using Procleave software ^39^. The sequences in bold indicate caspase 1, 3 and 6 putative cleavage sites with a probability score of > 0.7. The regions of MLB1 (top) and MLB2 (bottom) deletions are shown. **(D)** Huh7.5.1 cells were infected with indicated recombinant viruses of MLB1 (blue bars) and MLB2 (red bars) at an MOI 0.1 in triplicates, the virus was collected at 96 hours post infection, and titer was measured. Data are mean ± SEM. *p < 0.05, ns, non-significant using unpaired t-test test against wt MLB virus titer. **(E)** Analysis of CP expression in Huh7.5.1 cells infected with 2^nd^ passage of MLB1 and MLB2 mutant viruses (MOI 0.1). Cell lysates were harvested at 48 hpi and analyzed by western blotting with anti-CP and anti-tubulin antibodies. GNN is RdRp knock-out recombinant virus (GDD to GNN). **(F)** The effect of pan-caspase inhibitor z-VAD-fmk on capsid polyprotein processing during infection with classical human astrovirus 4 (HAstV4), MLB1 and MLB2 astroviruses. Caco2 cells were infected with HAstV4 and Huh7.5.1 cells were infected with MLB1 and MLB2 astroviruses at MOI 1 in the presence or absence of z-VAD-fmk. Cell lysates were harvested at indicated hpi and analyzed by western blotting with 8E7 antibody against HAstV CP (for HAstV4), anti-CP (for MLB) and anti-tubulin antibodies. The average inhibition of CP cleavage was quantified from 3 independent experiments.

### Generation of recombinant MLB viruses with deletions

To investigate the individual roles of truncations identified in the passaging experiments (Fig. 1D), we employed the RG system to create a set of recombinant viruses with the identified deletions (Fig. 4C). The titers of recombinant viruses were higher or comparable to the corresponding wt MLB1 and MLB2, with significantly lower titers only observed for MLB1-Δ42 (Fig. 4D). The analysis of infected cells was performed using a CP_NTD_-recognizing antibody, which was expected to recognize only N-terminal products of CP (Fig. 4E). Similarly to related HAstVs, the C-terminal domain of MLB1 and MLB2 CP contains putative caspase cleavage sites, which could lead to the programmed C-terminal cleavage of CP polyprotein. The predicted caspase cleavage sites were mapped for both MLB1 and MLB2 C-terminal domains of the capsid polyprotein (Fig. 4C). Consistent with experimental observations, the cleaved product of MLB2 CP corresponds to the size of CP in MLB2-Δ5^OF^ mutant suggesting caspase cleavage is taking place in this region (Fig. 4C, E). The shorter forms of MLB1 deletion mutants seem to locate downstream of predicted caspase cleavage sites and potentially affect the C-terminal cleavage of CP in different ways: Δ30 and Δ93 follow the wt-like processing of CP, Δ74^OF^ results in a shorter CP form, whereas Δ42 have both CP forms present. Notably, both Δ74^OF^ and Δ42 lack the lower cleavage product, presumably one of the C-terminal truncated forms (Fig. 4C-D). The integrity and stability of resulting mutant viruses over 3 passages were confirmed by RT-PCR and sequencing. To confirm the predicted caspase cleavage of CP, the Caco2 cells infected with classical human astrovirus 4 (HAstV4) and Huh7.5.1 cells infected with MLB1 and MLB2 were incubated in the absence and presence of pan-caspase inhibitor z-VAD-fmk. Consistent with the previously published results for classical HAstVs ^7^, both HAstV4 and MLB2 viruses resulted in the inhibition of CP cleavage in response to caspase inhibition (Fig. 4F). In contrast, MLB1 was not sensitive to the inhibition of caspase-mediated processes (Fig. 4F) suggesting that other cellular or viral proteases may be involved in the maturation of structural polyprotein of MLB1. These results support differences observed in the processing of CP that contains deletions in the C-terminal region.

### Creating MLB1 and MLB2 replicons to assess replication characteristics of truncated viruses

The deletions that occurred in the 3′ end of the genome could potentially affect the replication properties of the virus, considering the unequivocal importance of 3′ UTR in the replication of (+)ssRNA viruses. Multiple structured RNA elements are predicted in the 3′ UTR of MLB1 and MLB2 genomes, including 3 stable stem-loops mapped to the MLB1 deletion region (Fig. 5A). The 5-nucleotide deletion in MLB2 was not mapped to the structured RNA region, suggesting possible involvement of short- and/or long-range RNA interactions or other compensatory mechanisms including the processing of CP (Fig. 4E). All four deletions in MLB1 were mapped to the predicted stem-loop RNA structures (Fig. 5A) and could directly affect the RNA replication process.

**Figure 5.**
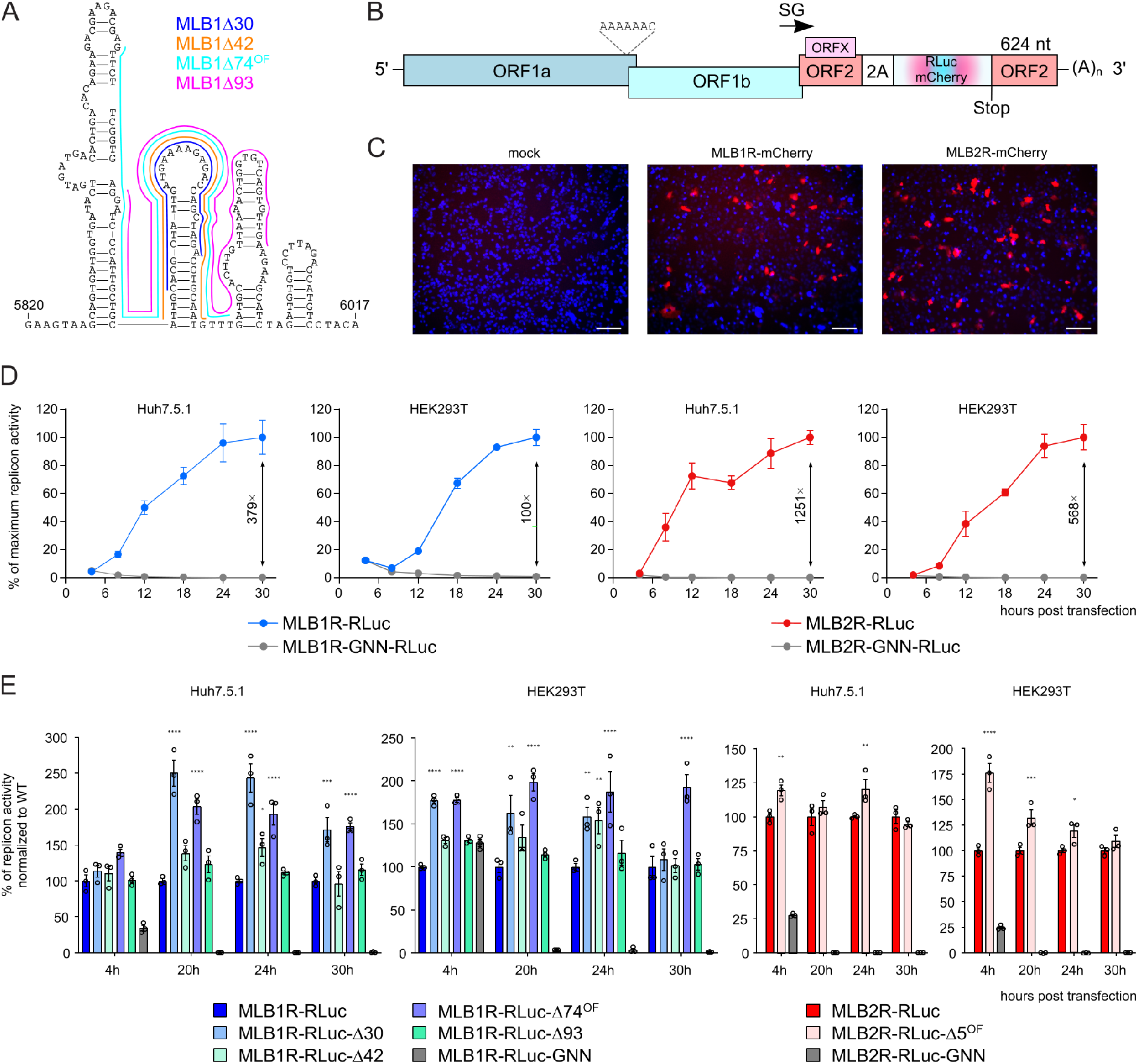
Creating MLB replicons to evaluate the importance of 3′ UTR. **(A)** The prediction of the RNA secondary structure and location of identified deletions in MLB1 3′ UTR. **(B)** Schematic of the MLB1 and MLB2 astrovirus replicons. The 2A-RLuc cassette is fused in the ORF2 followed by the stop codon and extended 3′ UTR. **(C)** Huh7.5.1 cells were transfected with MLB1 and MLB2 replicons expressing mCherry, incubated for 24 h, fixed and imaged, nuclei were counterstained with Hoechst (blue). Scale bars are 100 µm. **(D)** Relative MLB1 and MLB2 replicon luciferase activities were measured after RNA transfection of Huh7.5.1 or HEK293T cells. Values are normalized so that the mean wt replicon value at each time point is 100%. The replication fold difference between wt and GNN mutant replicon is provided for the final time point. **(E)** Relative MLB1 and MLB2 replicon luciferase activities were measured after RNA transfection of Huh7.5.1 or HEK293T cells. Data are mean ± SEM (n=3, ≥3 independent experiments, graphs D-E). *p < 0.05, **p < 0.01, ***p < 0.001, ****p < 0.0001 using two-way ANOVA test against wt replicon.

To evaluate this hypothesis and assess the importance of overlapping 3′ RNA structures, replicon systems for MLB1 and MLB2 were created. In replicon systems, the RNA replication is evaluated via a fluorescent or luminescent reporter gene that replaces the structural proteins. The preserved RNA elements contain intact SG promoter, 5′ and 3′ UTRs, functional RdRp and other components of the RNA replication machinery. This enables visualization and/or quantification of the SG reporter activity whilst avoiding the packaging step via deletion of the capsid region. Similar to a previously developed HAstV1-based replicon system ^5^, the genomes of MLB1 and MLB2 infectious clones were modified as demonstrated in Figure 5B. The 2A-cleavage mediated Renilla luciferase (RLuc) or mCherry reporters were used to quantify and visualize the protein expression from the SG promoter. The transfection of both MLB1R-mCherry and MLB2R-mCherry replicons resulted in the expression of reporter mCherry (Fig. 5C). The activity of MLB replicons expressing RLuc was monitored over time in two different cell lines (Fig. 5D). Strong replication was detected for MLB2 replicon in Huh7.5.1 (1251-fold) and HEK293T (568-fold) cells, when compared to replication-deficient but translation-competent GNN mutant replicon (with GDD catalytic RdRp motif changed to GNN). Lower replication levels were observed for MLB1 replicon in both Huh7.5.1 (379-fold) and HEK293T (100-fold) cells, which is consistent with slower growth kinetics observed for wt MLB1 virus (Fig. 3D-E).

To assess the replicon activity for MLB1 and MLB2 deletion mutants, all mutations (Fig. 1D) were transferred into corresponding RLuc replicon systems and activity was measured during the early (4 h) and later (20, 24 and 30 h) replication stages. A consistent 50-150% increase in replicon activity was observed for MLB1R-RLuc-Δ30 and MLB1R-RLuc-Δ74^OF^ mutants when compared to corresponding wt replicons in Huh7.5.1 cells (*p* < 0.001 for 20, 24 and 30 hpt). In HEK293T cells, MLB1R-RLuc-Δ74^OF^ was replicating above wt levels (*p* < 0.0001 for all time points) with other mutants having non-significant or less pronounced effects. A wt-like level was detected for MLB1R-RLuc-Δ42, MLB1R-RLuc-Δ93 and MLB2R-RLuc-Δ5^OF^ replicons at most time points (Fig. 5E). Taken together, these results suggest a beneficial or dispensable nature of identified 3′ RNA structures in the context of replicon system.

### MLB viruses with 3′ deletions are attenuated in iPSC-derived neurons

To assess the neurotropism of MLB viruses, we infected differentiated i^3^Neurons with wt and mutant recombinant MLB viruses at MOI 0.5 (Fig. 6A). The MLB1-Δ42 was not used due to lower titers in Huh7.5.1 cells (Fig. 4D). The infection with wt MLB2 reached 40.8% efficiency whereas MLB2-Δ5^OF^ showed weak signs of infection (4.8%), limited to single infected cells and not showing the spread of infection (Fig. 6B, D). The infection with wt MLB1 resulted in non-uniform distribution of infected cells with overall 8.6% CP-positive neurons. Conversely, the cells infected with MLB1-Δ30, MLB1-Δ74^OF^ and MLB1-Δ93 resulted in a decrease in infection efficiency (6.9%, 4.8% and 1.1% CP-positive cells, respectively) with MLB1-Δ93 being the most attenuated in primary neurons (Fig. 6C-D).

**Figure 6.**
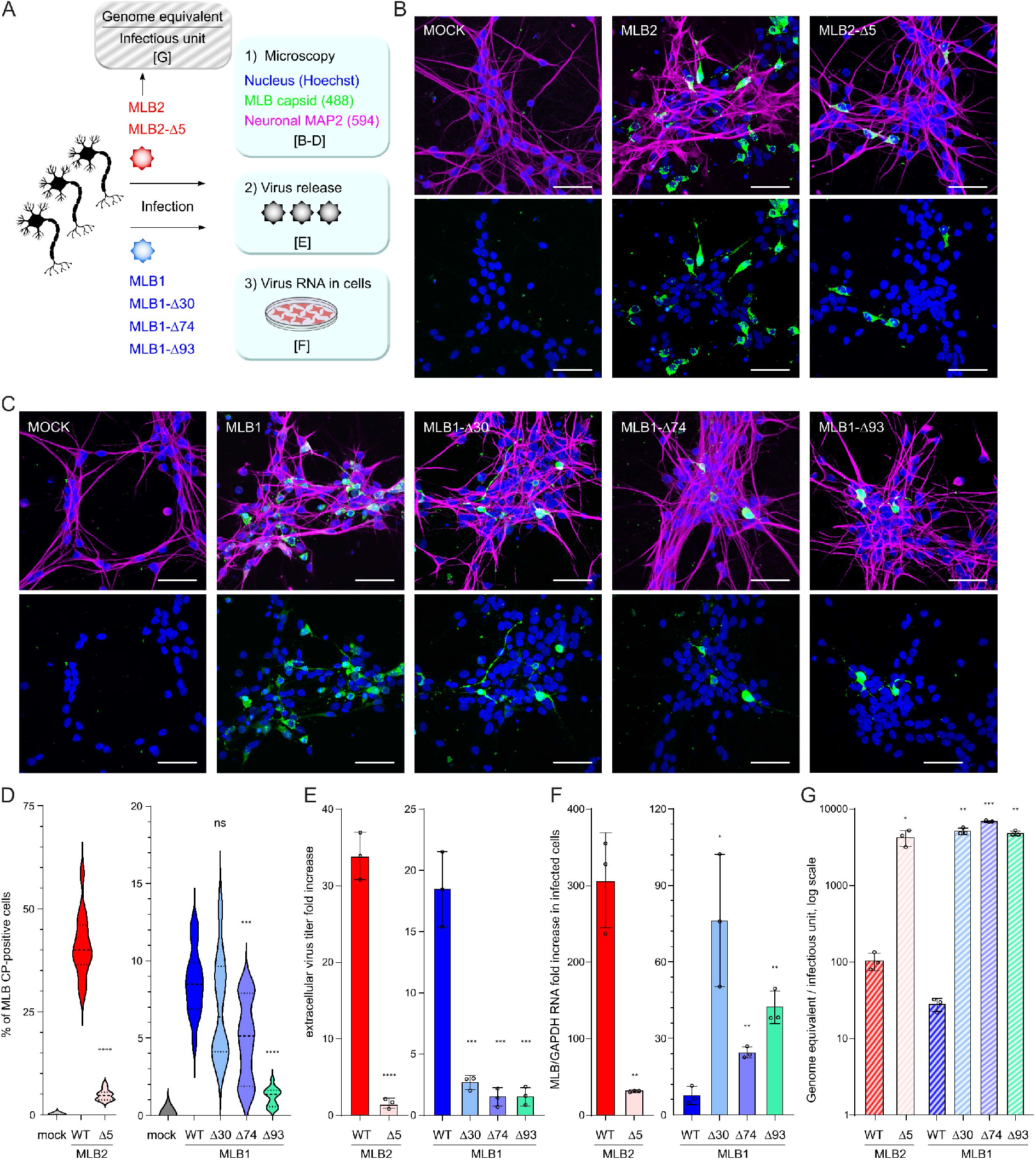
Infection of iPSC-derived i^3^Neurons with MLB astroviruses. **(A)** The schematic representation of the experiment where iPS cells were seeded on IBIDI (imaging) or 12-well (other analyses) plates, differentiated into mature i^3^Neurons, infected with indicated MLB1 and MLB2 recombinant viruses at MOI 0.5. **(B-C)** iPSC-derived neurons were infected with MLB2 (B) and MLB1 (C). Representative confocal images of fixed and permeabilized cells visualized for MLB CP (green) and MAP2 (magenta). Nuclei were stained with Hoechst (blue). Scale bars are 50 µm. **(D)** Approximately 15 images (150-250 cells per image) were analyzed for CP-positive cells. ***p < 0.001, ****p < 0.0001, ns non-significant, using two-tailed Mann-Whitney test against wt virus infection. **(E)** The virus release was measured by titration and normalized to the input virus titer. ***p < 0.001, ****p < 0.0001, ns non-significant, using two-tailed Mann-Whitney test against wt virus infection. **(F)** Intracellular RNA levels were quantified using qPCR. The virus-specific signal was normalized to GAPDH RNA and calculated as fold increase to the input RNA levels. **(G)** The genome equivalent to infection unit ratio was determined for MLB1 and MLB2 virus stocks by qPCR and virus titration, respectively. Data are mean ± SEM (n=3, ≥2 independent experiments, graphs E-G). *p < 0.05, **p < 0.01, ***p < 0.001, ****p < 0.0001 using two-way ANOVA test against wt infected samples.

Next, we examined the infectivity of the particles released from the infected neurons by titration on susceptible Huh7.5.1 cells. The analysis of virus release demonstrated that all viruses with deletions have significantly reduced production of infectious particles confirming that MLB astroviruses with deletions are strongly attenuated in neurons and result in decreased infectious virus release (Fig. 6E).

Next, we measured the levels of intracellular virus RNA in infected neurons. Consistent with lower infectivity (Fig. 6B, D, E), the virus-specific RNA transcripts were significantly reduced in MLB2-Δ5^OF^ (Fig. 6F). Surprisingly, the intracellular virus RNA levels were increased in MLB1 deletion mutants (Fig. 6F) despite lower CP-positive cells and reduced virus release (Fig. 6C-E). Coupled with similar observations in the replicon system (Fig. 5E), this suggests that RNA replication is strongly unbalanced in MLB1 deletion mutants and leads to decreased infectivity and particle release. These findings indicate that the 3′ region of MLB1 and MLB2 genome is required for establishing efficient infection in the neuronal cells, however, this is likely achieved in different ways for MLB1 and MLB2 mutants.

Finally, to examine if MLB1 and MLB2 mutants have altered specific infectivity, we analyzed the input Huh7.5.1-derived virus stocks for the presence of virus-specific RNA per number of infectious particles. As expected, for all deletion mutants this ratio was significantly altered when compared to the wild-type viruses (Fig. 6G).

Taken together, we developed a powerful platform to investigate neurotropic properties of MLB1 and MLB2 astroviruses and confirmed attenuation of the panel of recombinant viruses with specific 3′ deletions.

## Discussion

A novel MLB group of human astroviruses has gained increasing attention because of invading the non-gastrointestinal tract ^24,27,28^ and their zoonotic potential ^1,11^. However, the basic molecular biology of these viruses is still in its infancy because of the absence of genetically tractable *in vitro* infection models. Here we report a robust RG system for MLB1 and MLB2 genotypes of human astroviruses that allows the generation of recombinant viruses. We have used this to develop tools to efficiently propagate, visualize and quantify MLB-infected cells, and utilized this system to analyze MLB genomes with deletions in the 3′ region. Unlike previously reported astrovirus RG systems, MLB RG requires only one cell line for both rescue and virus passaging, enabling interrogation of early infection processes such as virus entry, uncoating and replication. This is particularly important for mutation-prone virus strains when prolonged infection results in substantial changes in the virus genome. As we observed for two recombinant MLB astroviruses, one of them (MLB2) had remarkable stability and no changes were detected after 10 serial passages. In contrast, MLB1 was less stable and had more variations, consistent with greater variability of previously reported sequences. This observation, coupled with the developed RG system highlights how investigating MLB represents an opportunity to study differential genomic stability in closely related viruses.

The processing of capsid polyprotein, virion assembly and maturation in MLB viruses is regulated by several cellular proteases, such as caspases. What is the role of C-terminal deletions in the context of ORF2-encoded polyprotein? The removal of the significant part of the acidic CP portion could affect CP localization, trafficking, particle formation and potential pro-viral roles that are yet to be characterized for astroviruses. In infected cells, we observed a major large polyprotein precursor and several smaller products (Fig. 2C, 3F), suggesting intracellular cleavage of CP. The analysis of media-derived samples revealed a prevalence of smaller CP cleavage products of about 55 kDa (Fig. 3G), indicating that CP is cleaved by additional unknown intracellular and/or extracellular proteases. The C-terminal cleavage of CP in MLB2 is likely to be regulated by cellular caspases (Fig. 4F), similar to classical human astroviruses ^7^. Despite the presence of the predicted cleavage sites (Fig. 4C), the processing of MLB1 capsid is not sensitive to cellular caspase inhibitors (Fig. 4F), resembling another neurotropic astrovirus, VA1 ^29^. In contrast to neurotropic VA1 and MLB groups of astroviruses, the infectivity of classical astroviruses strongly depends on exogenous trypsin activation ^2,29^. Since trypsin is a gut-specific enzyme, this may also explain the extra-gastrointestinal tract tropism of MLB astroviruses, including their ability to infect cell lines of different origins, such as Huh7 (hepatocarcinoma) and A549 (lung adenocarcinoma) ^2^. Taken together, the requirements for the proteolytic maturation of astrovirus capsid polyprotein represent a powerful strategy to control virus entry, infectivity and virion maturation, with a likely impact on cellular tropism and pathogenesis.

The deletions identified in the 3′ part of the MLB genomes can result in dual functional defect due to the overlap of structured RNA elements and ORF2 coding sequence. MLB1 and MLB2 have multiple differences in the key properties related to the functionality of the 3′ part of the genome: (i) RNA stem-loop structure was predicted for MLB1 (Fig. 5A), but not for MLB2, (ii) sensitivity to caspase inhibition was observed for MLB2, but not MLB1 (Fig. 4E), (iii) increased RNA replication in neurons and more significant increase of replicon activity was a hallmark for MLB1 deletion mutants (Fig. 5E, 6F). It was logical to expect that the 3′ attenuation in these two closely related MLB viruses could have differences in associated RNA and protein-related effects. All deletion mutants identified in MLB1 were mapped to the predicted structured stem-loops that could be beneficial for RdRp processivity in susceptible cells supporting active RNA replication. Enhanced genome replication was indeed observed for MLB1R-RLuc-Δ30 and -Δ74^OF^ replicons (Fig. 5E) and MLB1-Δ30, -Δ74^OF^ and -Δ93 viruses during infection in neurons (Fig. 6F), highlighting the importance of this region in the replication of MLB1 genome. The enhanced replication could have led to the imbalance of RNA replication-translation-packaging and resulted in lower infectious virus particle release and infectivity in neurons (Fig. 6C-E). In contrast, attenuation of MLB2 virus with a small 5-nucleotide out-of-frame deletion resulted in modest differences in replicon activity (Fig. 5E) and ten-fold decreased RNA levels in neuronal infection (Fig. 6F), suggesting that differences in cleaved C-terminal part of CP (Fig. 1D) could play a major role in attenuation of MLB2. This, however, does not exclude associated RNA defects (Fig. 5E, 6F-G) due to the overlapping nature of functional elements and tight association between RNA replication, translation, packaging and virus-host interactions.

The selection for deletion-prone viruses through serial passaging in highly susceptible cells is a well-known strategy for the generation of vaccine candidates ^22^. The viruses with deletions are usually more capable in cell culture and replicate to higher titers, but are highly attenuated in natural hosts, which makes them ideal candidates for live vaccines. Consistently, the deletion of the large portion of the acidic region in the context of the MLB genome could be well-tolerated in a highly susceptible Huh7.5.1 cell line but have a distinct role in the terminally differentiated neurons. To our knowledge, this is the first demonstration of attenuation strategy for neurotropic astroviruses, paving the way to the characterization of attenuation mechanisms and elucidation of the corresponding cell- and host-specific pathways. Similar vaccine candidates with deletions in accessory genome regions have been developed for various pathogenic viruses, including delta-6K and delta-5 (nsP3) mutants in Chikungunya virus ^30^, 30 nucleotide 3′ UTR deletion in Dengue virus ^31^, recently reported SARS-CoV-2 deletions in the multi-basic cleavage sites of spike protein ^32^, or combination of several known attenuation approaches that can provide a safer strategy to prevent reversion to virulence ^33^. Besides attenuation *in vivo*, trade-offs for enhanced cell culture replication may result in reduced particle stability ^34^. Taken together, the identification of the attenuation region in the MLB group of astroviruses brings us a step forward in understanding the mechanisms responsible for virulence in this group of viruses.

There is no animal model for human astroviruses reported so far to elucidate the attenuation mechanism in the context of systemic infection, but it would be interesting to introduce similar deletions in the genomes of closely related neuropathogenic animal astroviruses (Fig. 1A) to develop vaccine candidates against circulating astrovirus strains that cause outbreaks in animal farms ^14,35,36^.

In summary, we have developed a new RG system for MLB astroviruses and have exploited it to identify elements in the 3′ end of the genome that are dispensable for the replication in cell culture but attenuated in human neurons, thus developing our understanding of the molecular biology of MLB-group astroviruses and facilitating the development of therapeutics and vaccines. The identification of the regions responsible for the neurotropism in the MLB group of astroviruses represents first insights into molecular determinants of neuropathology of astroviruses, advances our understanding in mechanisms involved in this process and help identify virus strains with similar properties. This attenuation strategy can be further applied to other pathogenic human and animal astroviruses.

## Materials and Methods

### Cells

HEK293T cells (ATCC) were maintained at 37 °C in DMEM supplemented with 10% fetal bovine serum (FBS), 1 mM L-glutamine, and antibiotics. Huh7.5.1 (obtained from Apath, Brooklyn, NY) ^37^ and Caco-2 (ATCC) cells were maintained in the same media supplemented with non-essential amino acids (NEAA). All cells were tested mycoplasma negative throughout the work (MycoAlert® Mycoplasma Detection Kit, Lonza).

### Plasmids

For bacterial expression of CNP_NTD_, the relevant MLB1 CP-coding sequence corresponding to 61-396 aa in ORF2 was PCR amplified and inserted into the T7 promoter-based pExp-MBP-TEV-CHis expression plasmid with an N-terminal MBP fusion tag, followed by TEV protease cleavage site and C-terminal 8×His-tag (Fig. 1A).

To create RG clones for MLB1 and MLB2, the 5′ and 3′ terminal consensus sequences ^2^ were used to design specific primers to amplify MLB1 and MLB2 full-length genomes using Phusion™ High-Fidelity DNA polymerase (ThermoFisher Scientific). The amplified genomes of MLB1 and MLB2 were cloned into the T7 promoter-containing plasmid ^17^ using a single-step ligation independent cloning. Each 20 µl reaction was prepared using two PCR amplicons containing 15-20 nucleotide-long overlapping sequences mixed in equimolar proportions (50 ng for shorter product), 1× Buffer 2.1 (NEB), 2 µg BSA and 3 units of T4 DNA polymerase (NEB) and incubated at 20 °C for 30 minutes. The reaction was stopped by adding 1 µl of 20 mM dGTP, heated to 50 °C for 2.5 minutes, cooled down to room temperature for 20 minutes and used for the transformation of XL1 blue competent cells (Agilent). All identified mutations (Fig. 5A) were introduced in the MLB1 and MLB2 RG clones using site-directed mutagenesis. To create MLB1 and MLB2 replicon systems, both MLB RG clones were left intact up to the end of ORFX followed by a 2A sequence and a RLuc or mCherry sequence with a stop codon, followed by the last 624 nt of the virus genome and a 35 nt poly-A tail (Fig. 5B). All mutations were introduced into corresponding replicon plasmids using available restriction sites. The GenBank accession numbers for the pMLB1 and pMLB2 are ON398705 and ON398706, respectively.

All obtained plasmids were sequenced and annotated. The resulting RG and replicon plasmids were linearized with *Xho*I restriction enzyme prior to T7 transcription.

### Purification of His-tagged CP_NTD_ and generation of CP-specific antibody

The MLB1 CP_NTD_ protein was produced in Rosetta 2 (DE3) cells (Novagen) cultured in 2×YT media with overnight expression at 18 °C induced with 0.4 mM IPTG. The protein was purified first by immobilized metal affinity chromatography using PureCube Ni-NTA resin and then by affinity chromatography using amylose resin (NEB). N-terminal MBP fusion tag was removed by the cleavage with TEV protease (produced in-house). The MLB1 CP_NTD_ protein was further purified by heparin chromatography using HiTrap Heparin HP 5 ml column (Cytiva) and, finally, by size exclusion chromatography using a Superdex 200 16/600 column (Cytiva). Protein solution in 50 mM Na-phosphate pH 7.4, 300 mM NaCl, 5% glycerol was concentrated to 2 mg/ml and used for immunisation.

Antibody against CP_NTD_ was generated in rabbit using 5-dose 88-day immunisation protocol. Sera was used for CP_NTD_-specific affinity purification, followed by purification of specific IgG fraction (BioServUK Ltd).

### Recovery of MLB1 and MLB2 viruses from T7 RNAs

The linearized RG plasmids were used as templates to produce capped T7 RNA transcripts using T7 mMESSAGE mMACHINE™ T7 Transcription kit (Invitrogen) according to the manufacturer’s instructions. For a virus recovery, 10^7^ Huh7.5.1 cells were trypsinized, washed with PBS and electroporated with 20 µg T7 RNA in 800 µl PBS pulsed twice at 800 V and 25 µF using a Bio-Rad Gene Pulser Xcell™ electroporation system. The cell suspension was supplemented with 10% FBS-containing media and incubated at 37 °C. After 3 h of incubation and full cell attachment, the media was replaced with serum free media, and cells were incubated until appearance of CPE. To produce recombinant MLB1 virus stocks, electroporated cells were incubated for 72-96 h, freeze-thawed twice, filtered through 0.2 µm filter, supplemented with 5% glycerol and stored in small aliquots at -70 °C. For recombinant MLB2 virus stocks, electroporated cells were incubated for 48-72 h, the supernatant was clarified by filtration through 0.2 µm filter, supplemented with 5% glycerol and stored in small aliquots at -70 °C.

### Virus passaging, growth curves and titration

To passage recombinant MLB1 virus stocks, Huh7.5.1 cells were infected at an MOI 0.1 for 2 hours in serum-free media, then 5% FBS-containing media was added and incubated for 16-24 hours, then replaced with serum-free media and incubated for 72-96 h until the appearance of CPE, freeze-thawed twice, filtered through 0.2 µm filter and supplemented with 5% glycerol. To passage recombinant MLB2 virus stocks, Huh7.5.1 cells were infected at an MOI 0.1 for 2 hours in serum-free media, then 5% FBS-containing media was added and incubated for 16-24 hours, then replaced with serum-free media and incubated for 48-72 h until the appearance of CPE, the supernatant was clarified by filtration through 0.2 µm filter and supplemented with 5% glycerol. Virus RNA was isolated by Direct-zol RNA MicroPrep (Zymo research), followed by RT-PCR and Sanger sequencing of the virus genome.

To concentrate media-derived samples, the infected Huh7.5.1 cells were incubated for 96 hours, the media was collected, clarified using 0.2 µm filter and pelleted at 54,000 rpm (180,000 ×g) for 2 hours at 4 °C in a TLA-55 rotor in an Optima Max-XP tabletop ultracentrifuge (Beckman). The supernatant was removed, the pellet was resuspended in 50 mM Tris (pH 6.8) to obtain 20× concentrated sample and analyzed by SDS-PAGE followed by western blotting.

Multistep growth curves were performed using an MOI of 0.1. Individual infections were performed in triplicates. Both cell- and media-derived samples were collected in equal volume at 1, 6, 12, 24, 48, 72 and 96 h post infection and saved for virus quantification.

The immunofluorescence-based detection with anti-CP_NTD_ MLB1 antibody (1:300) was combined with infrared detection readout and automated LI-COR software-based quantification. Briefly, 24 h before infection, Huh7.5.1 cells were plated into 96 well plates (2-3×10^4^ cells/well). The 10-fold serial dilutions of virus stock in a round-bottom 96-well plate were prepared using serum-free media supplemented with 0.2% BSA. The cells were infected with prepared virus dilutions in duplicates, incubated for 24 hours, fixed with 4% paraformaldehyde (PFA), processed for immunofluorescence staining, scanned and counted as the number of capsid-positive signals. The titers were determined as infectious units per ml (IU/ml).

### SDS-PAGE and immunoblotting

Protein samples were analyzed using 8% SDS-PAGE. The resolved proteins were then transferred to 0.2 µm nitrocellulose membranes and blocked with 4% Marvel milk powder in PBS. Immunoblotting of MBL1 and MLB2 capsid protein was performed using anti-CP_NTD_ MLB1 antibody (custom-made rabbit polyclonal antibody, 1:3000). Anti-tubulin antibody (Abcam, ab6160, 1:1000) was used for the cellular target. Secondary antibodies (Licor IRDye 800 and 680, 1:3000) were used for IR-based detection. Immunoblots were imaged on a LI-COR ODYSSEY CLx imager and analyzed using Image Studio version 5.2.

### Inhibition of caspase cleavage in astrovirus-infected cells

Caco-2 cells were infected with HAstV4, Huh7.5.1 cells were infected with MLB1 and MLB2 astroviruses (MOI 5) in the presence or absence of 20 µM z-VAD-fmk (pan-caspase inhibitor, Promega). At indicated time post infection, cells were lysed and analyzed by immunoblotting using virus-specific antibodies. The capsid protein of HAstV4 was detected using astrovirus 8E7 antibody (Santa Cruz Biotechnology, sc-53559, 1:750).

### Analysis of CPE and fluorescent microscopy

For the analysis of virus-induced CPE, plasma membranes of infected cells were stained with Wheat Germ Agglutinin Alexa Fluor™ 488 Conjugate (WGA, ThermoFisher Scientific, 1:250) for 20 min, followed by fixation with 4% paraformaldehyde for 20 min, permeabilization with 0.05% Triton X-100 in PBS for 15 min and nuclei counter-staining with Hoechst (ThermoFisher Scientific) for 15 min. Cells were washed twice with PBS and imaged using EVOS fluorescence microscope. For the analysis of CP localization, the infected Huh7.5.1 cells were incubated for 24 hours, fixed and permeabilized as above. CP was detected using anti-CP_NTD_ MLB1 antibody followed by incubation with secondary antibody (Alexa Fluor 488-conjugated goat anti-rabbit, Thermo Fisher, A21441). Nuclei were counter-stained with Hoechst. The images are single plane images taken with a Leica SP5 Confocal Microscope using a water-immersion 63× objective.

### MLB replicon assay

Linearized replicon-encoding plasmids were utilized to produce T7 RNAs using mMESSAGE mMACHINE T7 Transcription kit, purified using Zymo RNA Clean & Concentrator kit and quantified by nanodrop. Huh7.5.1 and HEK293T cells were transfected in triplicate with Lipofectamine 2000 reagent (Invitrogen), using the reverse transfection protocol. Briefly, a mixture of 0.5 μl Lipofectamine 2000 and 0.5 μl OMRO (OptiMEM containing 40 units/ml RNaseOUT) was incubated for 5 minutes at room temperature before adding to a mixture of 100 ng T7 replicon RNA, 10 ng T7 Firefly luciferase-encoding RNA and 10 μl OMRO per transfection. After 20 min incubation at room temperature, 100 μl of the prewashed cells (10^5^ cells) were added to the transfection mixture, incubated at room temperature for 5 minutes, supplemented with 5% FBS, transferred to a 96 well plate and incubated for indicated time at 37°C (4-30 h). Replicon activity was calculated as the ratio of Renilla (subgenomic reporter) to Firefly (co-transfected loading control RNA, cap-dependent translation) using Dual Luciferase Stop & Glo Reporter Assay System (Promega) and normalized by the same ratio for the control wt replicon. Three independent experiments, each in triplicate, were performed to confirm the reproducibility of the results. To control for the transfection efficiency, MLB1 and MLB2 replicons encoding mCherry fluorescent protein were also transfected and visualized using EVOS fluorescent microscope (ThermoFisher Scientific).

### Growth and infection of iPSC-derived i^3^Neurons

i^3^Neuron stem cells were maintained at 37 °C in complete E8 medium (Gibco) on plates coated with Matrigel (Corning) diluted 1:50 in DMEM. Initial three-day differentiation was induced with DMEM supplemented with 1× N2 supplement (Thermo), 1× NEAA, 1× Glutamax and 2 μg/ml doxycycline. 10 μM Rock Inhibitor (Y-27632, Tocris) was added during initial plating of cells to be differentiated and cells were plated onto Matrigel coated plates. Differentiation medium was replaced daily and after three days of differentiation, partially differentiated neurons were re-plated into Cortical Neuron (CN) media in IBIDI wells coated with 100 μg/ml PLO. Cortical neuron media consisted of: Neurobasal Plus medium (Gibco) supplemented with 1× B27 supplement (Gibco), 10 ng/ml BDNF (Peprotech), 10 ng/ml NT-3 (Peptrotech), 1 μg/ml laminin (Gibco) and 1 μg/ml doxycycline. After initial replating, neurons were then maintained in CN media without doxycycline for a further 11 days until mature.

The fully differentiated neurons were infected with MLB astroviruses at an MOI 0.5 in neuronal media. After 2 hours, the virus inoculum was removed and replaced with 50% fresh – 50% conditioned media. After 48 hpi (MLB2) or 96 hpi (MLB1), the cells grown on IBIDI wells were fixed, permeabilized and stained with anti-CP_NTD_ MLB1 antibody followed by incubation with Alexa 488-conjugated secondary antibody, followed by the staining with antibody against neuronal marker MAP2 (Abcam, ab11268) and Alexa 597-conjugated secondary antibody. Nuclei were counter-stained with Hoechst. The confocal images are a projection of a z-stack images taken with a Leica SP5 Confocal Microscope using a water-immersion 63× objective. To analyze the percentage of CP-positive cells, the images were taken with EVOS fluorescence microscope. Approximately 15 images (150-250 cells per image) were analyzed. The infectivity of the particles released from the infected neurons was determined by titration on susceptible Huh7.5.1 cells as described above.

### Analysis of RNA levels in samples collected from infected neurons and virus stocks

Terminally differentiated neurons grown on 12-well plates (400,000 cells per well) were infected at MOI 0.5 and collected at 72 hpi. RNA was isolated by Direct-zol RNA MicroPrep (Zymo research), followed by quantitative reverse transcription-PCR (RT-qPCR) detection of virus (MLB1, MLB2) and cell-specific (GAPDH) RNAs. Results were normalized to the amount of GAPDH RNA in the same sample. Fold differences in RNA concentration were calculated using the 2−ΔΔCT method.

The absolute amount of MLB RNA in virus stock was determined by RT-qPCR. A 20 µl aliquot of each sample was mixed with 4×10^6^ plaque forming units (PFUs) of purified Sindbis virus (SINV) stock, which was used to control the quality of RNA isolation. RNA was extracted using the Qiagen QIAamp viral RNA mini kit. Reverse transcription was performed using the QuantiTect reverse transcription kit (Qiagen) using virus-specific reverse primers for SINV (GTTGAAGAATCCGCATTGCATGG), MLB1 (GTTGCACTGGCACCAGAGTC), MLB2 (GTGATAGTGAGGGATCTTCTGC). The known genome copy MLB standards were prepared using quantified purified T7 RNA transcripts of full length MLB genomes.

Quantitative PCR was performed in triplicate using SsoFast EvaGreen Supermix (Bio-Rad) in a ViiA 7 Real-time PCR system (Applied Biosystems) for 40 cycles with two steps per cycle. MLB and SINV-specific primers were used to quantify corresponding virus RNAs; the primer efficiency was within 95–105%. Quantitative PCR was performed in triplicate using SsoFast EvaGreen Supermix (Bio-Rad) in a ViiA 7 Real-time PCR system (Applied Biosystems) for 40 cycles with two steps per cycle.

qPCR primers: SINV-F (GAAACAATAGGAGTGATAGGCA), SINV-R (TGCATACCCCTCAGTCTTAGC), GAPDH-F (GCAAATTCCATGGCACCGT), GAPDH-R (TCGCCCCACTTGATTTTGG), MLB1-F (TTGCCAAGTGAGCCTTACAAAC), MLB1-R (TGCCATCAACAACTGGAAGCAC), MLB2-F (GATGTCTTTGGAATGTGGGTAAAG), MLB2-R (CTAGGTGCAGGTCCTTTCTTAG).

### Analysis of MLB1 and MLB2 sequences using the NCBI database

Available MLB1 and MLB2 complete and ORF2-specific genome sequences from NCBI deposited before January 2022 were identified and extracted from NCBI using Blastp. These sequences were then aligned using MUSCLE (https://www.ebi.ac.uk/Tools/msa/muscle/) and visualized using Bioedit software.

### RNAfold analysis of 3′ terminal MLB sequences

The secondary structures of the 3′ terminal MLB1 and MLB2 sequences were predicted by the RNAfold Server using default settings ^38^.

### Prediction of caspase putative cleavage sites

The putative caspase cleavage sites in C-terminal part of the MLB1 and MLB2 CP were analyzed by Procleave software ^39^ using default settings. Only predicted cleavage sites with probability score of >0.7 were considered.

### Statistical analyses

Data were graphed and analyzed using GraphPad Prism and MS Excel. Where appropriate, data were analyzed using two-way ANOVA test or two-tailed Mann-Whitney test. Significance values are shown as ****p<0.0001, ***p<0.001, **p<0.01, *p<0.05.

## Acknowledgments

This work was funded by a Sir Henry Dale Fellowship (220620/Z/20/Z) from the Wellcome Trust and the Royal Society and an Isaac Newton Trust/Wellcome Trust ISSF/University of Cambridge Joint Research Grant to V.L. J.E.D and A.S.N. are supported by a Wellcome Trust Senior Research Fellowship (219447/Z/19/Z) awarded to J.E.D. Authors thank Andrew Firth and Marko Hyvönen for the support at the beginning of this project.

## Author contributions

H.A. performed infection and transfection experiments, analyzed data and wrote the manuscript. A.L. designed and developed MLB detection tools. A.S.N., S.C.G. and J.E.D. developed a methodology and prepared iPSC-derived i^3^Neurons. E.B.W.F. performed computational analysis of publicly available MLB sequences. R.L.O. assisted with microscopy and protein analysis experiments. D.L.V. and S.G. performed experiments with clinical MLB isolates. V.L. developed MLB reverse genetics and replicons, wrote the manuscript, provided supervision and acquired funding.

